# Infant EEG microstate dynamics relate to fine-grained patterns of infant attention during naturalistic play with caregivers

**DOI:** 10.1101/2024.07.20.604427

**Authors:** Armen Bagdasarov, Sarah Markert, Michael S. Gaffrey

**Affiliations:** Department of Psychology & Neuroscience, Duke University, Reuben-Cooke Building, 417 Chapel Drive, Durham, NC, 27708, USA; Children’s Wisconsin, 9000 W. Wisconsin Avenue, Milwaukee, WI, 53226; Medical College of Wisconsin, Division of Pediatric Psychology and Developmental Medicine, Department of Pediatrics, 8701 Watertown Plank Road, Milwaukee, WI, 53226

**Author notes:** Corresponding author: Armen Bagdasarov **Email:**. **Author Contributions:** CRediT – AB: Conceptualization, Methodology, Software, Validation, Formal Analysis, Investigation, Data Curation, Writing - Original Draft, Writing - Review & Editing, Visualization, Project Administration. SM: Methodology, Investigation, Writing - Original Draft, Project Administration. MSG: Conceptualization, Methodology, Resources, Writing - Review & Editing, Supervision, Project Administration, Funding Acquisition. **Competing Interest Statement:** The authors declare no competing interest.

**Keywords:** infants, EEG, microstates, eye tracking, attention

## Abstract

As infants grow, they develop greater attentional control during interactions with others, shifting from patterns of attention primarily driven by caregivers (exogenous) to those that are also self-directed (endogenous). The ability to endogenously control attention during infancy is thought to reflect ongoing brain development and is influenced by patterns of joint attention between infant and caregiver. However, whether measures of infant attentional control and caregiver behavior during infant-caregiver interactions relate to patterns of infant brain activity is unknown and key for informing developmental models of attentional control. Using data from 43 infant-caregiver dyads, we quantified patterns of visual attention with dyadic, head-mounted eye tracking during infant-caregiver play and associated them with the duration of infant EEG microstate D/4 measured during rest. Importantly, microstate D/4 is a scalp potential topography thought to reflect the organization and function of attention-related brain networks. We found that microstate D/4 associated positively with infant-led joint attention rate but did not associate with caregiver-led joint attention rate, suggesting that infants’ initiatives in joint attention during play may be critical for the neurobiological development of attentional control. Further, we found that microstate D/4 associated negatively with infant attention shifts rate and positively with infant sustained attention duration, suggesting that increased stability of microstate D/4 may reflect maturation of attentional control and its underlying neural substrates. Together, our findings provide novel insights into how infant attentional control abilities and infant-caregiver visual behavior during play are associated with the spatial and temporal dynamics of infant brain activity.

**Significance Statement:** The ability to control attention endogenously during infancy has been associated with individual differences in future language skills and executive and self-regulatory functions. However, how the infant brain supports the development of attentional control is unknown. Using dyadic, head-mounted eye tracking during infant-caregiver play with toys and microstate analysis of infant EEG during resting state, we found associations between measures of attentional control and brain activity underlying attention-related brain networks. Specifically, our findings suggest that longer durations of EEG microstate D/4 may reflect the maturation of attention-related brain networks that support the development of infant attentional control. This is the first study to provide insight into the functional significance of microstate D/4 in relation to real-world behavior during infancy.

## Introduction

Infants (birth to 12 months of age) learn about the visual world by orienting and attending to the salient characteristics of their immediate, physical environment. Their caregivers encourage learning by creating sensitive and supportive interactions during play, which set the stage for the early development of attentional control. Attentional control is the ability to engage and disengage attention endogenously, or voluntarily, and flexibly (1, 2). During infancy, caregivers externally support much of their infant’s moment-to-moment attention. For example, caregivers present novel toys during play to attract attention and aid in soothing (3). However, infants also slowly develop the ability to exert control over their own attention during this time, and their rudimentary ability to do this during the first year of postnatal life has been associated with individual differences in future language skills (4) and executive and self-regulatory functions (5– 7). Recent research suggests that early brain development plays a key role in the emergence of attentional control during infancy (8, 9). It also suggests that identifying very early neurobiological correlates of attentional control are key to further informing developmental models of this skill and its place within the broader continuum of attention-related development. Nevertheless, few studies have investigated the neural correlates of attentional control during infancy, limiting our neurobiological understanding of this skill and its emergence during this important developmental period.

Electroencephalography (EEG) offers a direct, noninvasive, and low-cost measure of brain function and organization that can be collected from infants during various states of arousal or activity. Historically, researchers have used event-related potential (ERP) or frequency-based (i.e., power) EEG measures collected while infants watched stimuli (e.g., objects, faces) on a screen to investigate the neural correlates of sustained (10–14) or joint attention (see review by Bradley et al. (15)). More recently, Xie et al. (16) showed changes associated with global (i.e., network-based) EEG functional connectivity (i.e., weighted phase lag index and graph theory measures) that differed during – compared to in the absence of – sustained attention. In an effort to study the development of attention in a more naturalistic context, Phillips et al. (17) recorded EEG and videos of each partner’s face during infant-caregiver play. They found that endogenous oscillatory activity (i.e., theta power) did not increase before bouts of infant-led joint attention. Rather, infants showed neural activity associated with anticipatory processing (i.e., increased alpha suppression) when caregivers joined their attention, suggesting that infants perceive – at the neural level – when their caregivers respond to their initiations (17). Together, these studies have suggested that periods of sustained and joint attention are accompanied by intrinsically maintained tonic alertness (i.e., the brain maintains a steady level of vigilance or alertness), improved information processing efficiency, and distinct patterns of functional connectivity (7, 8). While informative for understanding the neurobiological underpinnings of developing attention skills during infancy, measures of attention in prior work have been limited by stationary eye trackers mounted to screens or in a fixed position in the lab, restricting the field of view and resulting attention-related information that can be captured from them.

Recent advancements in eye tracking technology – namely, the use of dyadic, head-mounted eye tracking (D-ET) in naturalistic contexts (19, 20) – have provided unique insights into how infant and caregiver behavior during play facilitate the development of attentional control. The fine-grained resolution at which D-ET captures egocentric, first-person visual information (e.g., 30 times per second) allows the quantification of not only what infants see but precisely how they allocate attention while freely moving their head during dynamically unfolding activities (e.g., play). One area that D-ET has proven very useful for generating new insights is the development of sustained, focused attention during infancy (21). When infants encounter a novel stimulus, they orient their attention to it, and they look at it for a period of time. The duration of sustained attention towards an object grows from infancy through early childhood and is thought to reflect the maturation of attention-related systems, transitioning from largely exogenous to endogenous control of attention. A growing body of D-ET research has provided unique insights into how caregivers and infants collaboratively shape the development of sustained attention. More specifically, recent D-ET work has suggested that infants use head and body movements to stabilize their visual attention to a specific target in the environment over distractors (i.e., closed-loop control hypothesis). Thus, as infants gain motor control and autonomy over their bodies, they actively manage their visual input (22, 23). Importantly, this cycle of attention and action is also influenced by caregiver behavior (24). For example, joint attention (i.e., when caregivers and infants coordinate their own visual attention with that of their partner to share a common point of reference (25)) has been shown with D-ET to increase the duration of infant sustained attention to objects during play (26), even after caregivers have looked away (27). Thus, with this work and that of others in mind, it has been suggested that changing patterns of infant attention during play reflect gradual developmental changes in how exogenous and endogenous attention systems interact rather than a distinct “developmental divide” (i.e., driven by one or the other at different ages). Nevertheless, how this developmental process unfolds across infancy and to what degree brain development supports and constrains it remains unknown. As a result, a better understanding of sustained attention-related brain-behavior associations during this key developmental period would likely provide novel insights into the emergence of endogenous attention as well as its interaction with other developmental processes to both shape and be shaped by infant and caregiver behavior (28).

As reviewed above, prior work strongly suggests that measures of attention acquired during infant-caregiver play are likely to be associated with infant brain function and organization measured using EEG. And, if present, they may provide new insights into how caregiver behavior and the infant brain support the development of attentional control in the context of everyday interactions. However, to date, no studies have examined the relationship between temporally and spatially fine-grained D-ET measures of attention during infant-caregiver play and reliable network-based EEG measures of intrinsic functional brain organization measured separately. Understanding whether increasingly endogenous control over attention during infancy is associated with intrinsic brain activity thought to reflect attention-related brain networks is a critical first step for investigating how changes in patterns of infant brain activity are shaped by – and/or shape – infant-caregiver interactions over development. In addition, given the challenges of collecting task-based EEG from infants and concerns about the reliability of some EEG task data at this age (29), establishing the presence (or absence) of associations between infant attention and patterns of infant functional brain organization that can be reliably measured is key for developing objective neurobiological markers of infant attentional control abilities. One promising and reliable network-based method for characterizing brain function and organization using EEG during infancy is microstate analysis. Microstates are patterns of scalp potential topographies that reflect very short periods (i.e., typically less than ∼150 ms) of synchronized neural activity (i.e., large-scale functional networks) evolving dynamically over time (30, 31). A small number of four to seven canonical microstates, each with a distinct spatial topography, have been replicated and consistently shown to explain the majority of topographic variance in the EEG signal recorded in the absence of external task demands (i.e., resting state, passive video-watching) in infants (32), children (33–35), and adults (see review by Michel & Koenig, (31)). Their temporal properties – global explained variance (GEV), duration, coverage, and occurrence (see Michel & Koenig, (31) for overview) – have been previously reported to be highly reliable even with small amounts of data (32, 36–40) and show unique variation in how their values associate with individual differences in behavior.

Critically, canonical microstate D/4, which shows a fronto-central topography, is thought to represent control and attention network function and organization. Specifically, EEG source localization and simultaneous EEG-functional magnetic resonance imaging (fMRI) studies of microstate D/4 have reported areas that overlap with control and attention networks in both children and adults (35, 41–43), including the dorsal frontoparietal, lateral frontoparietal, and midcingulo-insular networks. Recent work has also shown that the temporal properties of microstate D/4 are associated with error-related activity in 4-8-year-old children (35). And, further, that neural sources underlying microstate D/4 and an identified error-related microstate significantly overlap, suggesting that microstate D/4 is functionally associated with detecting salient events and a need for increased attentional control. Notably, studies using fMRI have suggested that control and attention networks can be identified during the first year of life, and that within-network integration continues to strengthen as between-network segregation emerges during the second year of life (8, 9). Since prior work strongly suggests that microstate D/4 is reflective of functional networks shown to support higher-order cognitive processes, including both top-down, goal-directed (i.e., dorsal attention network) and bottom-up, stimulus-driven (i.e., ventral attention network) control of attention (35, 44), it raises the possibility that microstate D/4 may similarly provide an objective measure of these networks during infancy and an opportunity to meaningfully investigate attention-related brain-behavior associations at this age.

The current, cross-sectional study investigated whether infant microstate D/4 (referred to as microstate 4 hereafter) duration during passive video-watching would associate with D-ET measures of attentional control as reported in prior work (i.e., infant-led and caregiver-led joint attention, the rate of infant attention shifts, and the average duration of infant sustained attention) during infant-caregiver free play with novel toys. Microstate duration is interpreted as reflecting an organized pattern of neural communication between brain regions that is stable for a given period of time. Developmentally, the duration of a given microstate potentially reflects the maturing functional stability – and/or dynamics – of its neural sources. As such, individual differences in the associations between microstate 4 duration and behavioral skills previously associated with these sources (i.e., control and attention networks) are likely to be present and reflective of developmental variation in these relationships. Importantly, microstate 4 duration has been shown to have high levels of stability and internal consistency during infancy (32), making it an ideal candidate for planned individual difference analyses. Our sample included 43, 5-10-month-old infants and their caregivers. Continuous EEG was recorded from infants while they watched up to 15 minutes of relaxing videos and D-ET was recorded during four minutes of infant-caregiver play with age-appropriate toys. Given that microstate 4 may represent control and attention-related network activity, we predicted that microstate 4 duration would associate with the rate of infant-led, but not caregiver-led joint attention, and the average duration of infant sustained attention (i.e., indicators of attentional control abilities). We also predicted that microstate 4 duration would associate with the rate of infant attention shifts, but in the opposite direction of the associations with joint and sustained attention (i.e., we expect attention shifts to decrease as attentional control abilities increase). Although associations between microstate 4 duration and behavior related to attention have been identified in previous work with adults, the patterns of findings are not clear, and no literature exists during infancy. Therefore, we did not make specific predictions about the directionality of each relationship (i.e., positive or negative) and considered these analyses to be exploratory.

## Results

Microstate Analysis. Five microstates were identified for the group of 43 infants (Fig. 1). Microstate 4 showed a fronto-central spatial topography, and its mean duration was approximately normally distributed (mean = 83.63; standard deviation = 6.44; range = 72.61-96.69; Fig. 1). Descriptive statistics for the other four microstates are available in SI Appendix, Table S1. The Spearman-Brown split-half reliability coefficient for the duration of microstate 4 was .86 (see SI Appendix for additional details), indicating excellent internal consistency as previously demonstrated in Bagdasarov et al. (32).

**Fig. 1.**
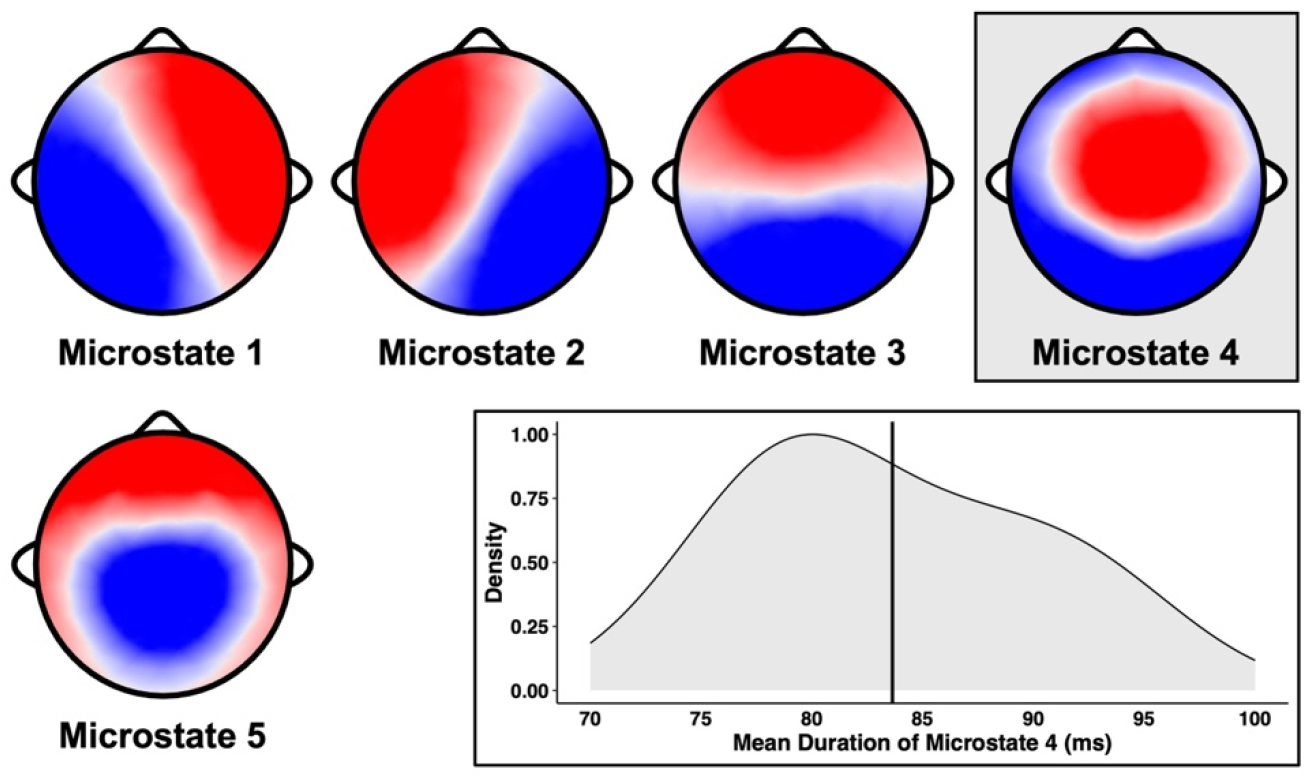
The five polarity-invariant infant EEG microstates. Together, they represented 73.43% of global explained variance at the group level and a mean of 57.76% (standard deviation = 2.69%; range = 53.51-64.50%) of global explained variance at the individual level after the backfitting procedure was performed. Microstate 4, which is highlighted in grey, represented 8.65% (standard deviation = 2.77%; range = 3.37-14.70%) of the global explained variance at the individual level after the backfitting procedure was performed. The distribution of the mean duration of microstate 4 is displayed below microstate 4. The black vertical line represents the mean of the distribution.

### Descriptive Statistics for D-ET Variables

*Infant-led* and caregiver-led joint attention rate. A paired samples *t-*test revealed that bouts of infant-led joint attention to toys (mean = 6.16 bouts per minute; standard deviation = 2.58; range = 1.16-12.64) occurred at a significantly higher rate than bouts of caregiver-led joint attention to toys (mean = 1.59 bouts per minute; standard deviation = 1.01; range = 0-3.91), t(42) = 11.77, p < .001 (Fig. 2A). Both the rate of infant-led and caregiver-led joint attention followed an approximately normal distribution, but the rate of infant-led joint attention showed much more variability across dyads.

**Fig. 2.**
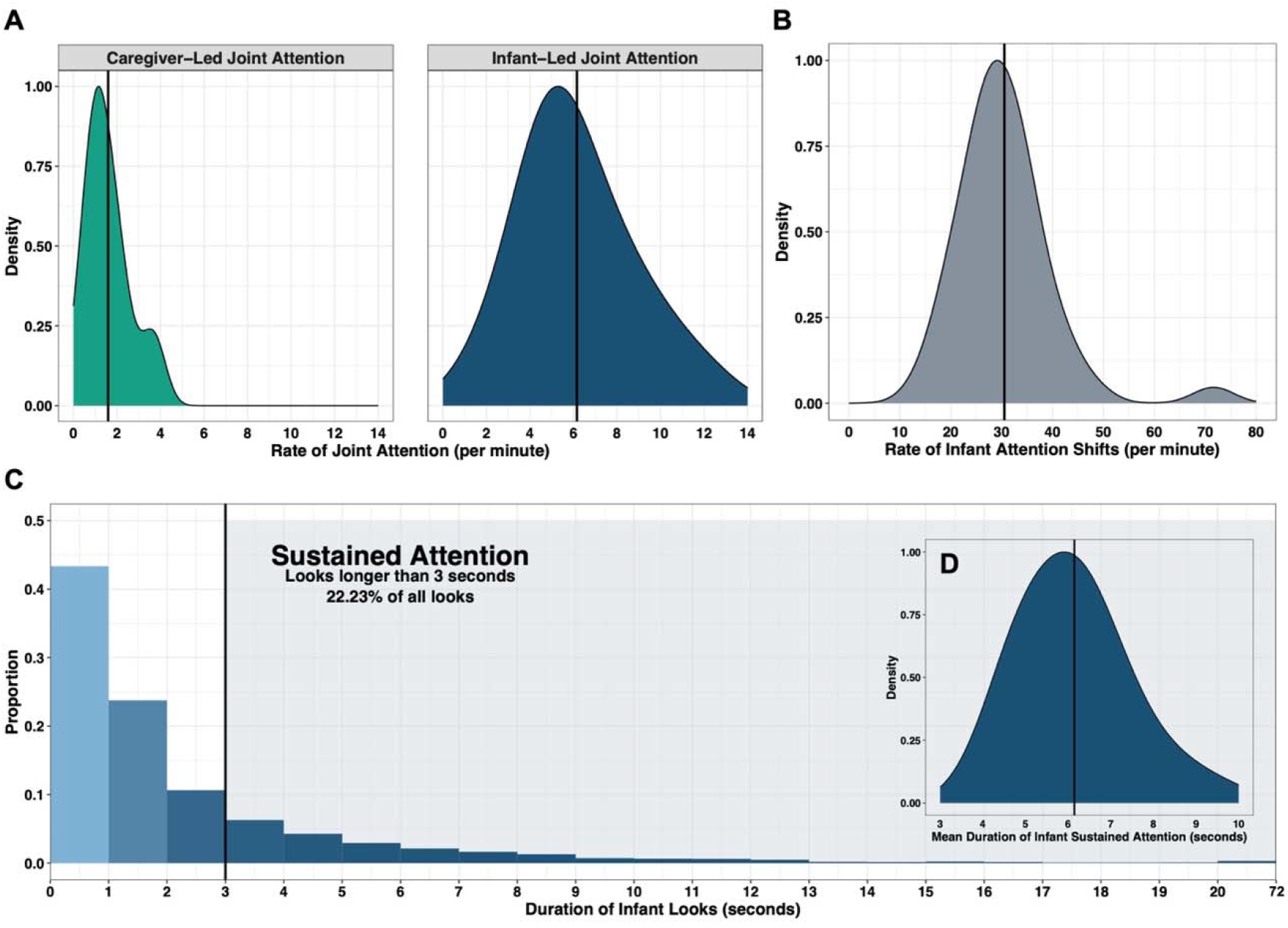
Distributions of D-ET variables. The black vertical lines displayed in A, B, and D represent the mean of the distribution. The black vertical line displayed in C represents the three-second mark. (A) Rate per minute of caregiver- and infant-led joint attention. (B) Rate per minute of infant attention shifts. (C) Distribution of all infant looks (n = 7,765). Sustained attention represented 22.23% of all infant looks (n = 1,751). A handful of looks (n = 24) were longer than 20 seconds in duration and, for visualization, they were lumped in the last bin representing 20 to 72 seconds. (D) Mean duration in seconds of infant sustained attention.

*Infant* attention shifts rate. The mean rate of infant attention shifts was 30.54 times per minute (standard deviation = 9.46; range = 14.48-71.63), which is similar to what has been previously reported in the literature (27). The mean rate of infant attention shifts followed an approximately normal distribution, except there was one outlier (i.e., one infant had a rate of 71.73 attention shifts per minute; Fig. 2B). When this outlier was removed, the mean rate of infant attention shifts was 29.57 (standard deviation = 7.04; range = 14.48-47.61).

*Infant* sustained attention duration. As observed in previous work (27), most infant looks to toys were brief in duration (mean = 2.38 seconds; standard deviation = 0.79; range = 1.13-4.47; Fig. 2C). Of all looks, 22.23% met the threshold for sustained attention. The mean duration of bouts of infant sustained attention followed an approximately normal distribution and as expected its mean across infants (mean = 6.15 seconds; standard deviation = 1.28; range = 3.99-9.79; Fig. 2D) exceeded the mean of the overall distribution of looks.

### Hierarchical Regression Analyses

All models included age and time of EEG data collection in the first step. The second step included one of the four D-ET variables: infant-led joint attention rate (model 1), caregiver-led joint attention rate (model 2), infant attention shifts rate (model 3), or infant sustained attention duration (model 4). For each model, regression tables with additional details are available in SI Appendix, Tables S2-S5. Multivariate outliers were found in the data for models 2 (n = 4), 3 (n = 2), and 4 (n = 2), and they were removed before performing each regression. See SI Appendix, Tables S6-S8 for models when these outliers were not removed.

*Model 1*. In step one, age and time of EEG data collection explained a significant proportion of the observed variation in microstate 4 duration, F(2, 40) = 6.07, adjusted *R*^2^ = .19, p = .005. In step two, the addition of infant-led joint attention rate was associated with a statistically significant increment to the proportion of the observed variation explained in microstate 4 duration, Δ adjusted *R*^2^ = .08, F(1, 39) = 5.08, *p* = .030. The final model containing all three predictors explained a significant proportion of the observed variation in microstate 4 duration, F(3, 39) = 6.15, adjusted *R*^2^ = .27, Bonferroni-corrected *p* = .006. Results indicated that a one standard deviation increase in infant-led joint attention rate associated with a .31 standard deviation increase in microstate 4 duration, all else held constant, t(39) = 2.25, *p* = .030 (Fig. 3A).

**Fig. 3.**
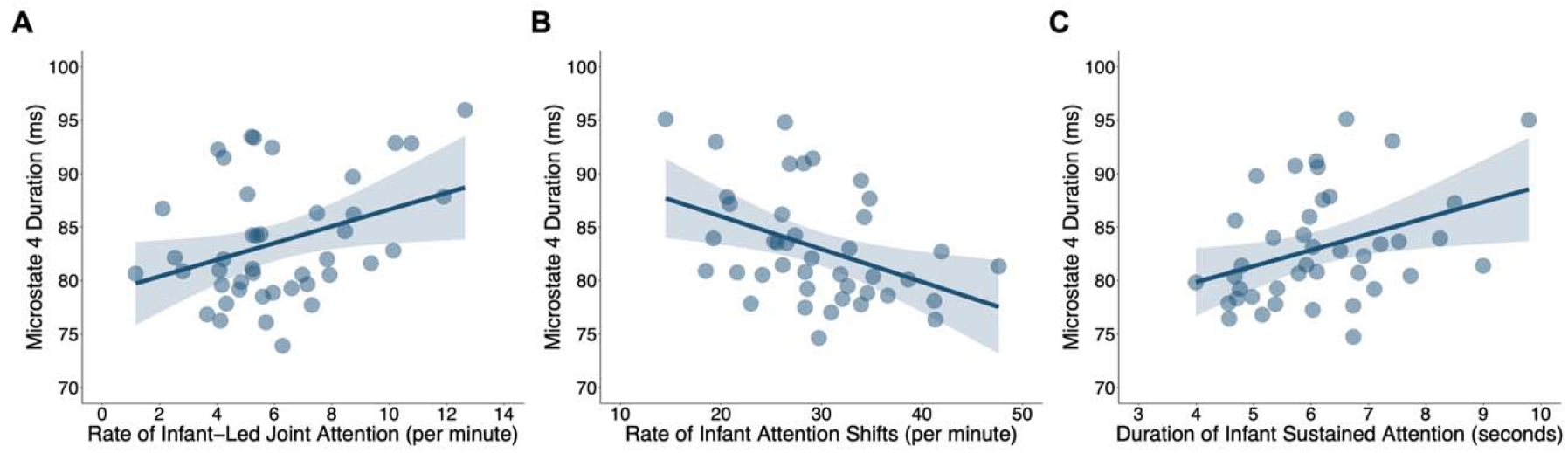
Partial residual plots for statistically significant regression models showing the effect of the independent variable on microstate 4 duration (ms) while holding age and time of EEG data collection constant. The shaded band represents the 95% confidence interval. Independent variables: (A) Rate per minute of infant-led joint attention (model 1). (B) Rate per minute of infant attention shifts (model 3). (C) Duration in seconds of infant sustained attention (model 4).

*Model 2*. In step one, age and time of EEG data collection explained a significant proportion of the observed variation in microstate 4 duration, *F*(2, 36) = 4.55, adjusted *R*^2^ = .16, *p* = .017. In step two, the addition of caregiver-led joint attention rate was not associated with a statistically significant increment to the proportion of the observed variation explained in microstate 4 duration, Δ adjusted *R*^2^ = .00, *F*(1, 35) = 0.99, *p* = .33. The final model containing all three predictors did not explain a significant proportion of the observed variation in microstate 4 duration, F(3, 35) = 3.36, adjusted *R*^2^ = .16, Bonferroni-corrected *p* = .118.

*Model 3*. In step one, age and time of EEG data collection explained a significant proportion of the observed variation in microstate 4 duration, F(2, 38) = 6.81, adjusted *R*^2^ = .23, *p* = .003. In step two, the addition of infant attention shifts rate was associated with a statistically significant increment to the proportion of the observed variation explained in microstate 4 duration, Δ adjusted *R*^2^ = .11, F(1, 37) = 7.64, *p* = .009. The final model containing all three predictors explained a significant proportion of the observed variation in microstate 4 duration, F(3, 37) = 7.88, adjusted *R*^2^ = .34, Bonferroni-corrected *p* = .001. Results indicated that a one standard deviation increase in infant attention shifts rate associated with a .36 standard deviation decrease in microstate 4 duration, all else held constant, t(37) = -2.76, *p* = .009 (Fig. 3B).

*Model 4*. In step one, age and time of EEG data collection explained a significant proportion of the observed variation in microstate 4 duration, F(2, 38) = 6.81, adjusted *R*^2^ = .23, p = .003. In step two, the addition of infant sustained attention duration was associated with a statistically significant increment to the proportion of the observed variation explained in microstate 4 duration, Δ adjusted *R*^2^ = .08, F(1, 37) = 5.75, *p* = .022. The final model containing all three predictors explained a significant proportion of the observed variation in microstate 4 duration, F(3, 37) = 7.02, adjusted *R*^2^ = .31, Bonferroni-corrected *p* = .003. Results indicated that a one standard deviation increase in infant sustained attention duration associated with a .32 standard deviation increase in microstate 4 duration, all else held constant, t(37) = 2.40, *p* = .022 (Fig. 3C).

## Discussion

We examined whether fine-grained D-ET measures of infant and caregiver attention during naturalistic infant-caregiver play associated the duration of infant EEG microstate 4 during resting state. We found that an increased rate of infant-led joint attention associated with an increased duration of microstate 4. However, there was no association between the rate of caregiver-led joint attention and the duration of microstate 4. Also, we found that an increased rate of infant attention shifts associated with a decreased duration of microstate 4. Lastly, an increased duration of infant sustained attention associated with an increased duration of microstate 4. Therefore, as predicted, the duration of microstate 4 associated with the rate of infant-led joint attention and duration of infant sustained attention, and in the opposite direction (positive) of its association with the rate of infant attention shifts (negative).

Our findings indicate that the frequency of infant-led joint attention bouts measured using D-ET varies substantially across infants and is associated with individual neurobiological differences measured at rest using EEG. Specifically, we showed that the rate of infant-led joint attention bouts positively associated with the duration of microstate 4 while caregiver-led ones did not. This may suggest that when infants are given space to drive their own attention toward an object, and that of their caregiver’s (i.e., a goal-directed behavior), they support the maturation of their own attention-related brain networks (i.e., infant self-organizes (45)). Interestingly, prior D-ET work has reported a positive association between the duration of infant-caregiver joint attention to an object and infant sustained attention to the same object after joint attention has ended (27). As a result, an alternative, though not mutually exclusive interpretation is that caregivers support the development of attention-related brain networks by creating a context that minimizes distractions and supports continued visual processing of salient objects (i.e., caregiver scaffolding (46)), though our study did not directly assess this. Nevertheless, the current results indicate that the frequency of infant-led interactions during play is likely a key factor shaping the maturation of brain networks associated with developing attentional control and is an area for future study (47).

Longer periods of sustained attention during play are an important marker of increasingly endogenous control over attention during infancy. Given this, the positive association between the durations of sustained attention and microstate 4 in the current study suggests that larger values of microstate 4 duration reflect the ongoing development of attention-related brain networks known to support increasingly mature attentional control (e.g., dorsal frontoparietal/orienting, lateral frontoparietal/control, and midcingulo-insular/executive networks). Importantly, as infants’ abilities in sustaining attention grow, they also spend less time shifting their attention between objects as they become better at resisting environmental distractions. In line with this, our finding that higher rates of infant attention shifts were associated with decreased duration of microstate 4 further supports the suggestion that larger values of microstate 4 duration reflect more mature control and attention-related brain network development. In fact, prior work has demonstrated that more efficient orienting (i.e., less frequent and unnecessary shifts of attention) is associated with increased activity of the executive network, which includes the anterior cingulate gyrus, anterior insula, basal ganglia, and parts of the prefrontal cortex (48, 49). This suggestion is also in line with prior adult studies showing increased temporal properties of microstate 4 during tasks requiring increased attentional demands (41, 50–52). As a result, the current findings indicate that the duration of microstate 4 may reflect ongoing maturation of attention-related brain function and organization during infancy, though longitudinal studies will be needed to fully address this possibility.

Our findings introduce the possibility that microstate 4 duration may serve as a measure of attention network function and organization in even younger infants (e.g., < 3 months of age). More specifically, microstate 4 has been identified in neonates during sleep (53). In addition, an adult-like frontoparietal network has been identified using fMRI during the same period (54). Together these prior findings suggest that microstate 4 may similarly represent functional organization of control and attention networks during very early infant development. However, prior studies have not investigated whether these neurobiological measures (e.g., microstate 4 duration) are associated with attentional control at these ages and/or later in development. Such information is likely critical for understanding whether temporal properties of microstate 4 such as duration can act as a useful neurobiological marker of very early developing attentional control and/or later emerging behaviors dependent upon the successful development of this skill (e.g., language, executive, and self-regulatory abilities). It is also likely to inform whether earlier emerging features of exogenous attention shape the later emerging endogenous attention system (i.e., Interactive Specialization (55)). Investigating this question and whether and how microstate 4 during infancy is predictive of later cognitive and behavioral outcomes during early childhood will require large samples to assess these longitudinal possibilities. Fortunately, multiple large-scale, longitudinal infant EEG studies such as HEALthy Brain and Child Development Study (56), Bucharest Early Intervention Project (57), Bangladesh Early Adversity Neuroimaging Study (58), Safe Passage Study (59), YOUth Cohort Study (60), Eurosibs Consortium (61), and Baby Siblings Research Consortium (62) are now underway and will provide important opportunities for doing this type of work and the development of novel analytical approaches in support of it (e.g., machine learning).

Our findings also contribute to the ongoing debate about the developmental timing of the emergence of attentional control during infancy by suggesting that it may begin to appear sooner than traditionally thought. As shown here, even during the first year of life, infants actively direct their own attention rather than responding solely to external cues. This may be accomplished through the integration of motor, control, and attention-related neural circuity supporting both exogenous and endogenous systems of attention (22, 23). To better understand the emergence of attentional control during infancy, a logical next step may therefore be to study its development in populations who are at-risk for altered exogenous attention, which emerges earlier in development (63). For example, a recent review and meta-analytic study found that infants born prematurely demonstrated a short-term advantage in visual-following during the neonatal period, but heightened risk for visual attention impairments cascading from more reflexive, exogenous (i.e., visual-following) to more complex, endogenous functions (i.e., visual recognition memory, sustained attention) during the first two years of postnatal life (64). As a result of these difficulties with visual attention in prematurely born infants, caregivers may subsequently change the way they interact with their infants (e.g., increased directedness or intrusiveness or less engagement during play) and, unintentionally, disrupt the natural development of attentional control (65). At the same time, infants may be less intrinsically motivated to actively explore and learn from their social environment. For example, infants with autism are likely to experience altered infant-caregiver interactions due to the disruptions in social engagement that characterize this disorder. As a result, they may fail to benefit from caregiver scaffolding of development and learning during this period and experience delays in skills dependent upon it, including language, executive, and self-regulatory functions (21, 65). Toward this end, and following models of developmental psychopathology, investigating the association between attentional control and microstate 4 duration in infants at risk for poor developmental outcomes is likely to provide a greater mechanistic understanding of how infant-caregiver interactions influence the neurodevelopmental biology underlying the early emergence of endogenous attentional control. Such research is now needed.

### Strengths and limitations

This is the first study to investigate the association between the temporal properties of EEG microstates and D-ET eye tracking measures of visual attention during infant-caregiver play, furthering our developmental understanding of the associations between infant-caregiver interactions, infant attention, and brain activity. Critically, although our study is a step toward measuring real-world infant behaviors and relating it to brain activity, infant-caregiver interactions during four minutes of play are unlikely to fully represent how infants and caregivers interact at home in their everyday environment. That is, we only captured a small snapshot of infant-caregiver behavior. Still, these simple interactions have been shown to have an enormous impact on learning over time (66). For example, joint attention is incrementally refined over time through experience over the course of early development (27, 67, 68). Further, we did not quantify other sensory and motor behaviors known to contribute to episodes of joint and sustained attention. Yet, prior work has shown that caregivers engage in different types of behaviors within joint attention beyond coordination of their eye gaze including talk and touch to support infant sustained attention (26, 69, 70). It will be important for future work to quantify all of these caregiver behaviors to build a more comprehensive understanding of how caregivers support of infant initiatives in joint attention associates with developing brain function and organization. However, given the time-consuming nature of coding D-ET data (i.e., it took our D-ET coders eight hours on average to code just the eye gaze of each dyads’ data), novel innovations in automated video-coding approaches (71, 72) are critical for this and now needed. Finally, given the lack of temporal alignment in our measures of brain and behavior (i.e., EEG and D-ET were not collected at the same time), we are not able to inform whether the duration of microstate 4 shapes and/or is shaped by our measures of attentional control. Still, microstates are considered to reflect scale-free patterns of neural activity (73) and may provide insight into how fast neural processes support behavioral states that unfold over longer time scales. The goal of our study was to explore brain-behavior associations as a first step toward developing causal and predictive models in the future.

## Conclusion

In conclusion, our findings indicate that features of infant attention during play are uniquely associated with specific spatial and temporal dynamics of electrical brain activity. These findings also further underscore how studying infant-caregiver interactions during play can lead to novel insights about the development of attentional control and the maturation of associated brain activity. Critically, this work employed a highly reliable approach for measuring infant brain activity (i.e., microstate analysis) and eye tracking measures that reflect attention in real-world contexts. As a result, and in consideration of future research, the current study highlights the potential for developing EEG microstate 4 as a neurobiological marker of attentional control development during infancy.

## Materials and Methods

### Participants

Infant-caregiver dyads were part of a larger study investigating the impact of bias and discrimination on prenatal and postnatal maternal health and infant development. All caregivers identified as female. Study details including recruitment and eligibility are available in Bagdasarov et al. (32). Sixty-five infant-caregiver dyads participated in one or two lab visits when infants were approximately 5-7-months and/or 8-10-months of age. Of these, 43 infant-caregiver dyads met the following inclusion criteria for the current study: (a) infants were required to have at least four minutes of preprocessed EEG data as recommended by previous work assessing the reliability of EEG microstates from infants (32), (b) infant-caregiver dyads were required to have at least 1.5 minutes of preprocessed D-ET data for coding, and (c) the EEG and D-ET data must have been collected on the same day. The mean age of infants was 8.31 months (standard deviation = 1.53; range = 5.98-10.42; see SI Appendix, Figure S1 for histogram) and 23 (53.49%) of them were biological males. Additional participant eligibility details are available in the SI Appendix, and participant demographics are available in SI Appendix, Table S9. All research was approved by the Duke University Health System Institutional Review Board and carried out in accordance with the Declaration of Helsinki. Caregivers provided informed consent and compensation was provided for their participation.

### EEG Data Collection, Preprocessing, and Microstate Analysis

Infants sat on their caregiver’s lap and watched up to 15 minutes of dynamic, relaxing videos with sound while 128-channel EEG (Electrical Geodesics, Eugene, OR) was recorded at 1000 hertz (Hz). Preprocessing was performed with EEGLAB (74) in MATLAB (MathWorks, Natick, MA). Details about EEG data collection and preprocessing are published in Bagdasarov et al. (32) and summarized in the SI Appendix. Quality control (QC) was performed (see SI Appendix, Table S10 for descriptive statistics) and participants whose data did not pass QC criteria (e.g., at least four minutes of preprocessed data) were excluded (see the SI Appendix for QC criteria). Each remaining participant’s data was cut to include only the first four minutes to reduce potential effects of varying data lengths on brain-behavior analyses. The preprocessed data were submitted to Cartool (75) and spatially filtered to remove topographic outliers and smooth topographies (76).

Microstate analysis using polarity-invariant k-means clustering (77) with resampling (32, 33, 35, 78) was performed in Cartool following the procedure published in Bagdasarov et al. (32) and summarized in the SI Appendix. An aggregate measure of six independent statistical criteria – the meta-criterion (41, 43) – determined the optimal number of microstates at both individual and group levels. Each time point of each participant’s preprocessed, spatially-filtered data was assigned a group-level microstate (i.e., backfitting procedure with a minimum spatial correlation of .50) and the average duration of microstate 4 was calculated for each participant. Importantly, choosing microstate 4 duration a priori significantly reduced the number of statistical tests performed.

### D-ET Data Collection and Coding

Caregivers and their infants sat across from each other at a small table and wore a head-mounted eye tracker built by Positive Science, LLC (Rochester, NY). Caregivers wore glasses while infants wore a comfortable hat, each with an infrared camera pointed to their right eye and a scene camera capturing their first-person view with 90° of visual angle (19). The D-ET calibration procedure is described in the SI Appendix. See Fig. 4 for D-ET setup.

D-ET was recorded at 30 Hz (i.e., 30 frames per second) while caregivers and their infants engaged in approximately four, one-minute periods of natural free play with one of two sets of three small, age-appropriate toys (e.g., rubber octopus, plastic plane). The order of sets was counterbalanced across trials (i.e., ABAB). This number of sets and toys was chosen to encourage infants to demonstrate different facets of their visual attention (e.g., orienting, shifting, and sustaining attention), and it is in line with previous work using a similar procedure (27, 69, 79). Caregivers were instructed to play as they naturally would at home for the duration of the four minutes. D-ET ended earlier than four minutes if the infant became fussy and/or cried. One experimenter operated the D-ET recording computer and monitored data collection in real time (Experimenter 1 in Fig. 4). Another experimenter switched the toy sets every minute and placed toys back onto the table if they fell off (Experimenter 2 in Fig. 4). A third experimenter sat directly behind the infant to prevent them from touching their D-ET hat (Experimenter 3 in Fig. 4). All experimenters were out of view of the infant during active data collection. Also, all experimenters and caregivers (but not infants) wore a face mask as part of the university’s COVID-19 health and safety precautions. For consistency, face masks were worn even after precautions were lifted.

A team of trained research assistants coded the exact location of the caregiver’s and infant’s gaze frame-by-frame as indicated by a crosshair in Noldus’ Observer XT (80). Descriptive codes are provided in the SI Appendix. D-ET data was coded only if at least 1.5 minutes of good data was provided by the infant-caregiver dyad. This was determined as data without infant fussiness and/or crying or issues with the corneal reflection which can interfere with D-ET accuracy (81). Infant-caregiver dyads had a mean of 4.27 minutes of D-ET data (standard deviation = 0.38; range = 3.10-5.11). The training procedure for coders is described in the SI Appendix. To ensure sustained reliability after training, 20% of all D-ET sessions of the larger study were coded twice (infant kappa mean = .84, standard deviation = .06; caregiver kappa mean = .84, standard deviation = .05; see the SI Appendix for details).

The coded data were read into R (82) and the following variables were quantified using custom scripts: (a) rate per minute of infant-led joint attention to toys (i.e., when infant looked at toy first), (b) rate per minute of caregiver-led joint attention to toys (i.e., when caregiver looked at toy first), (c) rate per minute of attention shifts that the infant made between anything in the environment, and (d) average duration in seconds of infant sustained attention to toys. A bout of joint attention was defined as the continuous alignment of caregiver and infant fixation within the region-of-interest for a single toy that lasted longer than 500 milliseconds but that could include looks briefer than 300 milliseconds elsewhere (27). A bout of sustained attention was defined as the continuous fixation within the region-of-interest for a single toy without any looks elsewhere for longer than three seconds (27). Additional details about how variables were quantified are available in the *SI Appendix*.

### Statistical Analysis

Four hierarchical regression models were run in R with microstate 4 duration as the dependent variable. The first step always included age in months and the time of EEG data collection (0 = morning, 1 = afternoon), which were both correlated with microstate 4 duration (*SI Appendix, Table S11*). The second step included one of the four independent variables of interest: infant-led joint attention rate (model 1), caregiver-led joint attention rate (model 2), infant attention shifts rate (model 3), or infant sustained attention duration (model 4). Model change statistics assessed whether the second step improved each model’s performance. For each model, multivariate outliers were identified by the Minimum Covariance Determinant and removed. All models met regression assumptions. Analyses were performed without removing outliers, too (*SI Appendix*). To minimize the Type 1 error rate, a Bonferroni-correction for four comparisons (i.e., the total number of models) was applied to the p value of each omnibus F test of each full model, which tested the overall significance of the complete set of predictors (*SI Appendix, Tables S12 and S13*).

## Data Availability

Raw and preprocessed EEG data are available in the Brain Imaging Data Structure (BIDS) format (85) on OpenNeuro (86) (OpenNeuro Accession Number: ds005343; https://openneuro.org/datasets/ds005343/). Data and scripts for statistical analyses as well as the coded eye tracking data are available on GitHub (https://github.com/gaffreylab/Infant-Microstates-and-Attention/). Video data are not available to maintain the privacy of participants. All other data and scripts are available upon reasonable request at the discretion of the corresponding author.

## Acknowledgments

This work was supported by the Charles Lafitte Foundation Program in Psychological and Neuroscience Research at Duke University (MSG). The first author (AB) was supported by the National Science Foundation Graduate Research Fellowship. We thank Ellen Gaffrey, Margaret O’Brien, Sua Cho, and Spandan Goel for their help with data collection and coding.

[Due to BIORXIV policy, this figure is not included in the preprint because it contains images of participants.]

**Fig. 4.** D-ET setup and example codes. “Other” (i.e., the box shaded in grey) represents all gaze points other than on one of the toys (e.g., partner’s face, table, etc.) for example/visualization purposes only. Participants shown here consented to using their images.

