## Supplementary Materials for "Infant EEG microstate dynamics relate to fine-grained patterns of infant attention during naturalistic play with caregivers"

### Supporting Information Text

**Split-half reliability of microstate 4 duration.** The Spearman-Brown split-half reliability coefficient for the duration of microstate 4 was calculated using the same procedure described in Bagdasarov et al. (1). Each participant's data was split into six, equal, 40-second segments, and backfitting was performed twice; once on even segments and once on odd segments, resulting in two sets of duration values. The Spearman-Brown split-half reliability coefficient was then calculated in R using the *splithalfr* package (2).

**Participant eligibility details.** Criterion 1: 54 of 65 (83.08%) infants had at least four minutes of preprocessed EEG data from one of their lab visits. Criterion 2: 60 of 65 (93.31%) infants had at least 1.5 minutes of preprocessed D-ET data for coding from one of their lab visits. When combined, 48 of 65 (73.85%) infants met Criteria 1 and 2. Criterion 3: 43 of 48 (89.58%) infants had EEG and D-ET data collected on the same day. Data collection for the final set of 43 participants occurred between March 2022 and October 2023.

Given the longitudinal design of the parent study, some participants had EEG and ET data at both the six- and nine-month study visits. In this case, the visit closest to when the infant was exactly 6.00 or 9.00 months of age was selected so that each infant was only represented once in the current study's cross-sectional analyses. However, if the selected visit had less than four minutes of preprocessed ET data and the other visit had more data, then the other visit was selected to maximize the reliability of ET measures.

**EEG data collection and preprocessing.** Details about EEG data collection and preprocessing are published in Bagdasarov et al. (1), and are available on https://github.com/gaffreylab/EEG-Microstate-Analysis-Tutorial. Briefly, twenty-four outer ring channels that often contain a large amount of artifact in data from infants were removed. Data were downsampled to 250 Hz and bandpass filtered 1-20 Hz. Periods of inattention were manually rejected based on session notes. Bad channels were identified, removed, and interpolated if they were flat for more than five seconds, contained more than four standard deviations of line noise relative to all other channels, or correlated at less than .80 to surrounding channels. Data were re-referenced to the average. Independent component analysis (ICA) with principal component analysis dimensionality reduction to 50 components was performed on a copy of the data that was cleaned with Artifact Subspace Reconstruction (ASR; bust criterion of 20 to remove, but not reconstruct artifacts) (3, 4). The ICLabel plugin was used to flag components that had a probability of at least .70 of being related to noise from eye, muscle, or heart activity, or line or channel noise (5). Flagged components were removed from the full-length data which was not cleaned with ASR. Data were segmented into one-second epochs and removed in two steps using the TBT plugin (6). First, data containing residual eye-related artifacts were removed if any one of six frontal channels contained amplitudes > 150 or < -150 µV. Then, epochs were removed if at least 10 channels contained amplitudes > 150 or < -150 µV, joint probabilities above three standard deviations, or a 100 µV maximum-minimum amplitude difference between samples. If less than 10 channels met rejection criteria, the epoch was not removed, but the channels were interpolated for that epoch only. The data were re-referenced to the average again.

**EEG data quality control (QC).** Participants were only included in analyses if they met the following QC criteria:

1. Bad Channels
    - Number of bad channels must not exceed 15 (approximately 15% of 105 total channels).
    - Visualization of bad channels must not show clusters of bad channels.
2. Artifact Subspace Reconstruction (ASR)
    - Data length after ASR must exceed 60 seconds.
3. Independent Component Analysis (ICA)
    - Visualization of decomposition must appear "normal" and "appropriate."

- Visualization of flagged components must appear artifact-related.
- Retained variance after removal of flagged components must exceed 50%.

4. Channel Power Spectra
    - Visualization of channel power spectra must appear "normal" and "appropriate."
5. Data Duration
    - At least four minutes or 240 seconds of preprocessed data must be available.

**EEG microstate analysis.** The following is as described in Bagdasarov et al. (1) and available on https://github.com/gaffreylab/EEG-Microstate-Analysis-Tutorial.

Stage 1: Individual-Level Clustering

At the individual-level (i.e., for each participant's data), topographies at global field power (GFP) peaks representing timepoints of the highest signal-to-noise ratio were extracted (7). Fifty epochs, each composed of 8000 random subsamples of the extracted topographies and representing 99.9% of the participant's data, were submitted to a polarity-invariant modified *k*-means cluster analysis (8), which was set to repeat 50 times and identify 1-12 clusters of topographies for each epoch. The resampling approach is thought to improve the reliability of *k*-means clustering and has been used in recent work (1, 9–11). The meta-criterion – an aggregate measure of six independent criteria (12, 13) – determined the optimal number of clusters for each epoch, resulting in *k* optimal clusters for each of 50 epochs or *k**50 topographies for each participant.

Stage 2: Group-Level Clustering

At the group-level (i.e., for data from the group of participants), the 50 epochs of *k* optimal clusters from each participant were combined, resulting in 50**k* topographies for each of 43 participants or 50**k**43 topographies for the group. One-hundred epochs, each composed of 700 random subsample of these topographies and representing 99.9% of the group's 50**k**43 topographies were submitted to a polarity-invariant modified k-means cluster analysis, which was set to repeat 100 times and identify 1-15 clusters of topographies for each epoch. The meta-criterion determined the optimal number of clusters for each epoch, resulting in *k* optimal clusters for each of 100 epochs or *k**100 topographies. These topographies were combined and submitted to a final *k*-means cluster analysis with the same parameters, and the meta-criterion was used as guidance from which we selected the optimal number of clusters based on resting-state topographies observed in prior work; now, the group-level microstates.

Stage 3: Backfitting

The last stage of microstate analysis – backfitting – was performed on each participant's preprocessed, spatially-filtered data, and resulted in rich temporal information characterizing each group-level microstate for each participant. First, each participant's data was normalized by the median of GFP to account for individual differences in scalp potential due to varying skull conductivity. Then, each time point of each participant's data was labeled with one group-level microstate; the one that was most spatially correlated with the topography at that time point (i.e., winner-take-all approach). The polarity of topographies was ignored when calculating the correlation and the minimum correlation for time points assigned to a microstate was .50. After backfitting, temporal smoothing was applied (window half-size of 32 ms and Besag factor of 10; (8)), and improbably small segments were removed, such that segments smaller than 32 ms were divided in half with the first half added to the preceding segment and the second half added to the proceeding segment. Lastly, the duration values of each microstate were calculated for each participant.

**D-ET calibration and processing.** To calibrate the infant's eye tracker, an experimenter displayed a series of laser points on the table to capture the infant's attention. To calibrate the caregiver's eye tracker, caregivers were instructed to keep their heads still and visually track a point in the center of a black-and-white card provided by Positive Science, LLC (Rochester, NY). These calibration procedures were performed at the onset of the paradigm and as needed between trials if any shifts in equipment occurred (e.g., if the infant touched the eye tracking equipment). At the end of the play session, calibration points were collected again.

All videos were saved after each session and uploaded to Yarbus eye tracking software from Positive Science, LLC (Rochester, NY). Prerecorded eye and scene videos were used to complete a simple calibration process, which involved identifying calibration points from each participant's first-person camera view (i.e., navigating to calibration frames and clicking on fixation points). A multi-calibration procedure was followed if equipment shifted during play, which involved detecting the frame at which the shift occurred and identifying the calibration points collected following the shift to re-calibrate the data. Processed videos – displaying a small circle indicating the participant’s gaze point throughout the session – were rendered and saved for each participant.

**List of D-ET codes.**

- Blue Toy
- Green Toy
- Red Toy
- Partner’s Face
- Partner’s Body
- Partner’s Hand
- Other
- Extraneous Eye Movement
- Pupil or Corneal Reflection Detection Difficulties
- Toy Switch Present

**D-ET training and reliability.** Research assistants were trained to code eye tracking data using Noldus’ Observer XT. Processed videos overlaid with a small circle indicating caregiver and infant gaze points were uploaded to Observer XT. Coders navigated through each video frame-by-frame and identified whether the gaze point of each participant was on a toy (red, blue, yellow), partner's body, partner's hand, partner's face, or other location (e.g., the table, an experimenter). Coders also indicated frames at which the gaze point could not be determined (i.e., due to poor eye image, obstruction of eye image, participant interference with equipment, or infant crying). This approach is consistent with previous studies using D-ET technology (14). Coders reached reliability when they completed three consecutive videos with at least 85% of frames in agreement with a master coder. A second coder independently coded 20% of all videos part of the larger study for reliability upkeep and to ensure inter-coder agreement of over 85% of frames. *Note: Percent agreement was used for training and sustained reliability. Kappa values are reported in the main manuscript.*

**Tables S1 to S13**

**Table S1.** Descriptive statistics for each microstate's duration.

| | Mean | Standard Deviation | Min | Max |
|---|---|---|---|---|
| Microstate 1 | 78.19 | 3.89 | 71.91 | 88.56 |
| Microstate 2 | 78.37 | 3.52 | 69.67 | 85.51 |
| Microstate 3 | 101.76 | 5.15 | 91.48 | 115.89 |
| Microstate 4 | 83.63 | 6.44 | 72.61 | 96.69 |
| Microstate 5 | 82.18 | 4.83 | 71.67 | 94.56 |

All values are in milliseconds (ms).

**Table S2.** Model 1 regression results with outliers identified and removed (*n* = 0 outliers).

| Variable | *B* | 95% CI for *B* | | *SE B* | β | $R^2$ | $\Delta R^2$ |
|---|---|---|---|---|---|---|---|
| | | LL | UL | | | | |
| Step 1 | | | | | | .19** | .19** |
| Constant | 98.10*** | 88.08 | 108.12 | 4.96 | | | |
| Age | -1.54* | -2.72 | -0.36 | 0.59 | -0.37* | | |
| Time | -3.76* | -7.35 | -0.16 | 1.78 | -0.29* | | |
| Step 2 | | | | | | .27** | .08* |
| Constant | 96.70*** | 87.07 | 106.34 | 4.76 | | | |
| Age | -1.95** | -3.14 | -0.77 | 0.59 | -0.46** | | |
| Time | -3.79* | -7.22 | -0.36 | 1.69 | -0.30* | | |
| Infant-Led JA Rate | 0.78* | 0.08 | 1.49 | 0.35 | 0.31* | | |

CI = confidence interval; LL = lower limit; UL = upper limit; JA = joint attention. $R^2$ values were adjusted for the number of predictors. Full model *p* values were Bonferroni-corrected for four comparisons (i.e., significance asterisk next to the $R^2$ value in Step 2 represents the corrected *p* value).

Full Model: $F(3, 39) = 6.15$, adjusted $R^2 = .27$, original $p = .002$, corrected $p = .006$.

Infant-Led JA Rate: $t(39) = 2.25$, $p = .030$.

$^{+}p < .10$. $^{*}p < .05$. $^{**}p < .01$. $^{***}p < .001$.

**Table S3.** Model 2 regression results with outliers identified and removed (*n* = 4 outliers).

| Variable | *B* | 95% CI for *B* | | *SE B* | β | $R^2$ | $\Delta R^2$ |
|---|---|---|---|---|---|---|---|
| | | LL | UL | | | | |
| Step 1 | | | | | | .16* | .16* |
| Constant | 95.51*** | 85.17 | 105.85 | 5.10 | | | |
| Age | -1.22* | -2.44 | -0.01 | 0.60 | -0.31* | | |
| Time | -3.84* | -7.57 | -0.11 | 1.84 | -0.31* | | |
| Step 2 | | | | | | .16 | .00 |
| Constant | 95.42*** | 85.06 | 105.77 | 5.10 | | | |
| Age | -1.43* | -2.72 | -0.15 | 0.63 | -0.36* | | |
| Time | $-3.53^{+}$ | -7.32 | 0.26 | 1.87 | $-0.29^{+}$ | | |
| Caregiver-Led JA Rate | 1.25 | -1.31 | 3.82 | 1.26 | 0.16 | | |

CI = confidence interval; LL = lower limit; UL = upper limit; JA = joint attention. $R^2$ values were adjusted for the number of predictors. Full model *p* values were Bonferroni-corrected for four comparisons (i.e., significance asterisk next to the $R^2$ value in Step 2 represents the corrected *p* value).

Full Model: $F(3, 35) = 3.36$, adjusted $R^2 = .16$, original $p = .030$, corrected $p = .118$.

Caregiver-Led JA Rate: $t(35) = 0.99$, $p = .327$.

$^{+}p < .10$. $^{*}p < .05$. $^{**}p < .01$. $^{***}p < .001$.

**Table S4.** Model 3 regression results with outliers identified and removed (*n* = 2 outliers).

| Variable | *B* | 95% CI for *B* | | *SE B* | β | $R^2$ | $\Delta R^2$ |
|---|---|---|---|---|---|---|---|
| | | LL | UL | | | | |
| Step 1 | | | | | | .23** | .23** |
| Constant | 93.46*** | 83.56 | 103.36 | 4.89 | | | |
| Age | -0.98⁺ | -2.15 | 0.20 | 0.58 | -0.24⁺ | | |
| Time | -5.15** | -8.64 | -1.66 | 1.73 | -0.42** | | |
| Step 2 | | | | | | .34** | .11** |
| Constant | 102.67*** | 91.31 | 114.03 | 5.61 | | | |
| Age | -0.99⁺ | -2.07 | 0.10 | 0.53 | -0.24⁺ | | |
| Time | -5.39** | -8.61 | -2.16 | 1.59 | -0.44** | | |
| Attention Shifts Rate | -0.31** | -0.53 | -0.08 | 0.11 | -0.36** | | |

CI = confidence interval; LL = lower limit; UL = upper limit. $R^2$ values were adjusted for the number of predictors. Full model *p* values were Bonferroni-corrected for four comparisons (i.e., significance asterisk next to the $R^2$ value in Step 2 represents the corrected *p* value).
Full Model: $F(3, 37) = 7.88$, adjusted $R^2 = .34$, original $p < .001$, corrected $p = .001$.
Attention Shifts Rate: $t(37) = -2.76$, $p = .009$.
$^+p < .10$. $^*p < .05$. $^{**}p < .01$. $^{***}p < .001$.

**Table S5.** Model 4 regression results with outliers identified and removed ($n$ = 2 outliers).

| Variable | $B$ | 95% CI for $B$ | | $SE$ $B$ | β | $R^2$ | $\Delta R^2$ |
|---|---|---|---|---|---|---|---|
| | | LL | UL | | | | |
| Step 1 | | | | | | .23** | .23** |
| Constant | 93.46*** | 83.56 | 103.36 | 4.89 | | | |
| Age | -0.98⁺ | -2.15 | 0.20 | 0.58 | -0.24⁺ | | |
| Time | -5.15** | -8.64 | -1.66 | 1.73 | -0.42** | | |
| Step 2 | | | | | | .31** | .08* |
| Constant | 83.35*** | 70.68 | 96.01 | 6.25 | | | |
| Age | -0.85 | -1.96 | 0.26 | 0.55 | -0.21 | | |
| Time | -5.66** | -8.98 | -2.34 | 1.64 | -0.46** | | |
| SA Duration | 1.50* | 0.23 | 2.77 | 0.63 | 0.32* | | |

CI = confidence interval; LL = lower limit; UL = upper limit; SA = sustained attention. $R^2$ values were adjusted for the number of predictors. Full model $p$ values were Bonferroni-corrected for four comparisons (i.e., significance asterisk next to the $R^2$ value in Step 2 represents the corrected $p$ value).

Full Model: $F(3, 37) = 7.02$, adjusted $R^2 = .31$, original $p < .001$, corrected $p = .003$.

SA Duration: $t(37) = 1.50$, $p = .022$.

$^+p < .10$. $^*p < .05$. $^{**}p < .01$. $^{***}p < .001$.

**Table S6.** Model 2 regression results with outliers not removed (outliers kept in model).

| Variable | *B* | 95% CI for *B* | | *SE B* | β | $R^2$ | $\Delta R^2$ |
|---|---|---|---|---|---|---|---|
| | | LL | UL | | | | |
| Step 1 | | | | | | .19** | .19** |
| Constant | 98.10*** | 88.08 | 108.12 | 4.96 | | | |
| Age | -1.54* | -2.72 | -0.36 | 0.59 | -0.37* | | |
| Time | -3.76* | -7.35 | -0.16 | 1.78 | -0.29* | | |
| Step 2 | | | | | | .19* | .00 |
| Constant | 97.42*** | 87.23 | 107.61 | 5.04 | | | |
| Age | -1.60** | -2.80 | -0.41 | 0.59 | -0.38** | | |
| Time | -3.79* | -7.40 | -0.18 | 1.79 | -0.30* | | |
| Caregiver-Led JA Rate | 0.76 | -1.05 | 2.57 | 0.90 | 0.12 | | |

CI = confidence interval; LL = lower limit; UL = upper limit; JA = joint attention. $R^2$ values were adjusted for the number of predictors. Full model *p* values were Bonferroni-corrected for four comparisons (i.e., significance asterisk next to the $R^2$ value in Step 2 represents the corrected *p* value).
Full Model: $F(3, 39)$ = X.XX, adjusted $R^2 = .19$, original $p = .011$, corrected $p = .043$.
Caregiver-Led JA Rate: $t(39) = 0.85$, $p = .402$.
$^+p < .10$. $^*p < .05$. $^{**}p < .01$. $^{***}p < .001$.

**Table S7.** Model 3 regression results with outliers not removed (outliers kept in model).

| Variable | *B* | 95% CI for *B* | | *SE B* | β | $R^2$ | $\Delta R^2$ |
|---|---|---|---|---|---|---|---|
| | | LL | UL | | | | |
| Step 1 | | | | | | .19** | .19** |
| Constant | 98.10*** | 88.08 | 108.12 | 4.96 | | | |
| Age | -1.54* | -2.72 | -0.36 | 0.59 | -0.37* | | |
| Time | -3.76* | -7.35 | -0.16 | 1.78 | -0.29* | | |
| Step 2 | | | | | | .18⁺ | .01 |
| Constant | 100.11*** | 87.37 | 112.84 | 6.30 | | | |
| Age | -1.60* | -2.81 | -0.38 | 0.60 | -0.38* | | |
| Time | -3.65* | -7.31 | -0.004 | 1.81 | -0.29* | | |
| Attention Shifts Rate | -0.05 | -0.25 | 0.15 | 0.10 | -0.08 | | |

CI = confidence interval; LL = lower limit; UL = upper limit; JA = joint attention. $R^2$ values were adjusted for the number of predictors. Full model *p* values were Bonferroni-corrected for four comparisons (i.e., significance asterisk next to the $R^2$ value in Step 2 represents the corrected *p* value).
Full Model: $F(3, 39) = 4.06$, adjusted $R^2 = .18$, original $p = .013$, corrected $p = .053$.
Attention Shifts Rate: $t(39) = -0.53$, $p = .602$.
$^+p < .10$. $^*p < .05$. $^{**}p < .01$. $^{***}p < .001$.

**Table S8.** Model 4 regression results with outliers not removed (outliers kept in model).

| Variable | *B* | 95% CI for *B* | | *SE B* | β | $R^2$ | $\Delta R^2$ |
|---|---|---|---|---|---|---|---|
| | | LL | UL | | | | |
| Step 1 | | | | | | .19** | .19** |
| Constant | 98.10*** | 88.08 | 108.12 | 4.96 | | | |
| Age | -1.54* | -2.72 | -0.36 | 0.59 | -0.37* | | |
| Time | -3.76* | -7.35 | -0.16 | 1.78 | -0.29* | | |
| Step 2 | | | | | | .23* | .04$^+$ |
| Constant | 90.44*** | 77.30 | 103.58 | 6.50 | | | |
| Age | -1.49* | -2.65 | -0.34 | 0.57 | -0.35* | | |
| Time | -4.04* | -7.56 | -0.52 | 1.74 | -0.32* | | |
| SA Duration | 1.20$^+$ | -0.18 | 2.58 | 0.68 | 0.24$^+$ | | |

CI = confidence interval; LL = lower limit; UL = upper limit; JA = joint attention. $R^2$ values were adjusted for the number of predictors. Full model *p* values were Bonferroni-corrected for four comparisons (i.e., significance asterisk next to the $R^2$ value in Step 2 represents the corrected *p* value).
Full Model: $F(3, 39) = 5.29$, adjusted $R^2 = .23$, original $p = .004$, corrected $p = .015$.
SA Duration: $t(39) = 1.76$, $p = .086$.
$^+p < .10$. $^*p < .05$. $^{**}p < .01$. $^{***}p < .001$.

**Table S9.** Participant demographics.

| | Mean (SD) | Range |
|---|---|---|
| Age (months) | 8.31 (1.53) | 5.98-10.42 |
| | *n* | Percent (%) |
| Biological Sex | | |
| Males | 23 | 53.49% |
| Females | 20 | 46.51% |
| Race | | |
| White | 35 | 81.40% |
| Multiracial | 4 | 9.30% |
| Black or African American | 3 | 6.98% |
| Other | 1 | 2.33% |
| Asian | 0 | 0% |
| Native Hawaiian or Other Pacific Islander | 0 | 0% |
| Ethnicity | | |
| Not Hispanic or Latino | 38 | 88.37% |
| Hispanic or Latino | 5 | 11.63% |
| Maternal Education | | |
| Some Grade School | 0 | 0% |
| Completed Grade School | 0 | 0% |
| Some High School | 1 | 2.33% |
| High School Diploma | 1 | 2.33% |
| Some College or 2-Year Degree | 1 | 2.33% |
| 4-Year College Degree | 16 | 37.21% |
| Some School Beyond College | 1 | 2.33% |
| Professional or Graduate Degree | 23 | 53.49% |

All demographic variables were obtained at the time of study enrollment, except for age, which represents infants' ages at the time of their EEG and D-ET session. $n = 43$.

**Table S10.** Descriptive statistics of EEG data quality control metrics.

| | Number of Channels Removed and Interpolated | File Length in Seconds After ASR | Number of Independent Components Rejected | Percent Variance Retained After Removal of Independent Components | File Length in Seconds After Preprocessing |
|---|---|---|---|---|---|
| Mean | 4.42 | 480.29 | 4.09 | 89.21 | 436.51 |
| SD | 2.37 | 118.73 | 2.03 | 5.00 | 110.24 |
| Minimum | 0 | 286.49 | 1 | 74.77 | 272 |
| Maximum | 9 | 766.96 | 8 | 98.29 | 752 |

ASR = Artifact Subspace Reconstruction. SD = standard deviation. *n* = 43.

**Table S11.** Pearson's correlations between microstate 4 duration and age, biological sex, time of EEG data collection, and duration of D-ET.

| | Microstate 4 Duration | |
|---|---|---|
| | Correlation, *r* | *p* value |
| Age | -.38 | .011 |
| Biological Sex | -.14 | .379 |
| Time of EEG Data Collection | -.32 | .039 |
| Duration of D-ET | -.08 | .602 |

EEG data collection occurred in the morning (9:00am-11:59am) for 24 participants and in the afternoon (12:00pm-4:00pm) for 19 participants.

**Table S12.** Original and corrected model *p* values (outliers identified and removed).

| Original *p* value | Corrected *p* value |
|---|---|
| .002 | .006 |
| .030 | .118 |
| < .001 | .001 |
| < .001 | .003 |

A Bonferroni-correction for four comparisons (i.e., the total number of models) was applied to the *p* value of each omnibus *F* test of each full model, which tested the overall significance of the complete set of predictors.

**Table S13.** Original and corrected model *p* values (outliers kept in model).

| Original *p* value | Corrected *p* value |
|---|---|
| .002 | .006 |
| .011 | .043 |
| .013 | .053 |
| .004 | .015 |

A Bonferroni-correction for four comparisons (i.e., the total number of models) was applied to the *p* value of each omnibus *F* test of each full model, which tested the overall significance of the complete set of predictors.

**Figure S1**

**Figure S1.** Histogram of participant ages.

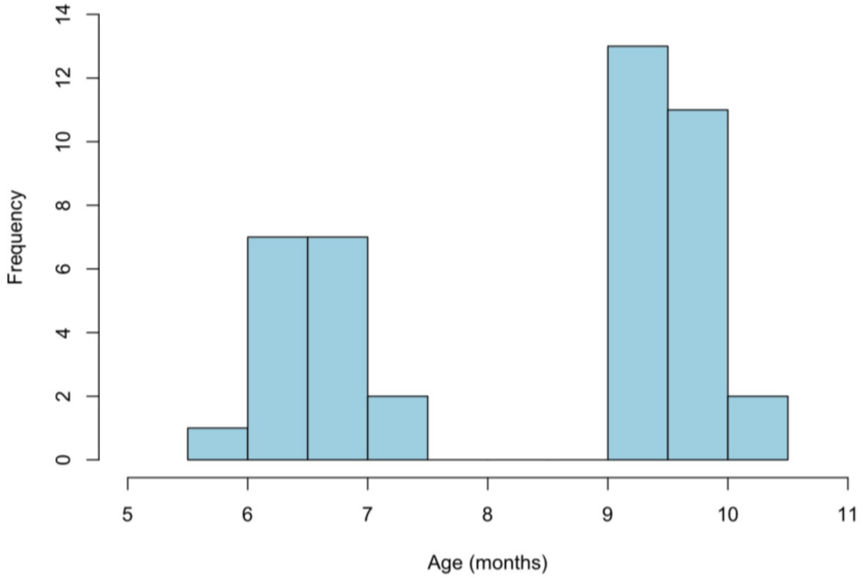

Infant-caregiver dyads participated in one or two lab visits when infants were approximately 5-7-months and/or 8-10-months of age. The final sample included 17 infants who were in the 5-7-months group and 26 infants who were in the 8-10-months group (total *n* = 43).
